# Genetic heterogeneity in depressive symptoms following the death of a spouse: Polygenic score analysis of the US Health and Retirement Study

**DOI:** 10.1101/065847

**Authors:** Benjamin W. Domingue, Hexuan Liu, Aysu Okbay, Daniel W. Belsky

**Affiliations:** Stanford University Graduate School of Education; School of Criminal Justice at the University of Cincinnati; Department of Complex Trait Genetics, Vrije Universiteit, Center for Neurogenomics and Cognitive Research, Amsterdam, the Netherlands; Erasmus University Rotterdam Institute for Behavior and Biology, Rotterdam, the Netherlands; Duke University School of Medicine, Department of Medicine, Division of Geriatrics, Duke Univeristy Social Science Research Institute, Duke University Center for the Study of Aging and Human Development

**Author notes:** The corresponding authors can be contacted at and.

**Keywords:** Stressful life events, depression, polygenic score, genome-wide association study, diathesis-stress

## Abstract

Experience of stressful life events is associated with risk of depression. Yet many exposed individuals do not become depressed. A controversial hypothesis is that genetic factors influence vulnerability to depression following stress. This hypothesis is most commonly tested with a “diathesis-stress” model, in which genes confer excess vulnerability. We tested an alternative model, in which genes may buffer against the depressogenic effects of life stress. We measured the hypothesized genetic buffer using a polygenic score derived from a published genome-wide association study (GWAS) of subjective wellbeing. We tested if married older adults who had higher polygenic scores were less vulnerable to depressive symptoms following the death of their spouse as compared to age-peers who had also lost their spouse and who had lower polygenic scores. We analyzed data from N=9,453 non-Hispanic white adults in the Health and Retirement Study (HRS), a population-representative longitudinal study of older adults in the United States. HRS adults with higher wellbeing polygenic scores experienced fewer depressive symptoms during follow-up. Those who survived death of their spouses during follow-up (n=1,829) experienced a sharp increase in depressive symptoms following the death and returned toward baseline over the following two years. Having a higher polygenic score buffered against increased depressive symptoms following a spouse's death. Effects were small and clinical relevance is uncertain, although polygenic score analyses may provide clues to behavioral pathways that can serve as therapeutic targets. Future studies of gene-environment interplay in depression may benefit from focus on genetics discovered for putative protective factors.

## INTRODUCTION

Experience of stressful life events is associated with risk of depression (1,2). Yet many individuals exposed to stressful life events do not become depressed. A controversial hypothesis is that genetic differences between individuals modify the influence of stressful life events on risk of developing mental health conditions, including depression (3). This “diathesis-stress” hypothesis is supported by family-based genetic studies that find individuals with familial liability to depression are more vulnerable to developing depression following stress exposure (4,5). Molecular genetic evidence for the diathesis-stress hypothesis from candidate-gene studies is contested, partly because of concerns about the candidate-gene approach (6). Genome-wide association studies (GWAS) offer opportunities to move beyond old arguments about candidate genes (7). GWAS discoveries for depression have yielded mixed results in tests of diathesis-stress models (8–10). Here, we test proof of concept for using an alternative GWAS phenotype to develop a genetic measures of susceptibility to depression following stress. Instead of a diathesis, such as genetic liability to depression, we focused on a potential measure of psychological robustness to stress, the genetics of subjective wellbeing.

Subjective wellbeing represents a dimension of so-called “positive psychology,” and may reflect the opposite end of an affective continuum from depression (11,12). Like depression, subjective wellbeing is heritable (13,14). A portion of this heritability is thought to reflect genetic influences shared with depression (15,16). Theory predicts increased subjective wellbeing should buffer against deleterious consequences of stressful life events (17,18).

We conducted a polygenic score study to test if genetic predisposition to greater subjective wellbeing buffered against risk of depression following experience of a stressful life event, the death of one's spouse. We studied depressive symptoms in a cohort of older adults and their spouses followed longitudinally as part of the US Health and Retirement Study. We combined whole-genome single-nucleotide polymorphism (SNP) data already generated for the cohort and results from GWAS of subjective wellbeing in over 100,000 adults by the Social Science Genetic Association Consortium (16) to calculate cohort members’ subjective wellbeing polygenic scores. Analysis tested if higher subjective-wellbeing polygenic score buffered survivors’ risk of developing depressive symptoms following the death of their spouse.

## MATERIALS AND METHODS

### Sample

The Health and Retirement Study (HRS) is a longitudinal survey of a representative sample of Americans over the age of 50 and their spouses initiated in 1992 (19,20). HRS is administered biennially and includes over 26,000 persons in 17,000 households. Respondents are interviewed about income and wealth, work and retirement, family and social connections, use of health services, and physical and mental health. A full description of the HRS is provided online at http://hrsonline.isr.umich.edu/index.php.

We linked HRS survey data from the “Rand Fat Files” (21) with genome-wide single nucleotide polymorphism (SNP) data downloaded from US National Institutes of Health Database of Genotypes and Phenotypes (dbGaP). HRS SNP genotyping was conducted by the NIH Center for Inherited Disease Research using DNA extracted from saliva collected during face-to-face interviews in HRS respondents’ homes in 2006 and 2008 and Illumina HumanOmni 2.5 Quad BeadChip arrays. SNPs missing in >5% of samples, with minor allele frequency <1%, or not in Hardy-Weinberg equilibrium (p<0.001) were removed. The final genetic database included approximately 1.7M SNPs. HRS SNP data are described in detail at http://hrsonline.isr.umich.edu/index.php?p=xxgen1.

Our analysis is based on 9,453 HRS respondents (42% male) who self-reported non-Hispanic white race/ethnicity. Respondents were born between 1905-1975 and were 59.5 years old on average at first interview (IQR=53-66). Included respondents participated in a median of 8 (IQR 6-10) follow-up assessments, yielding 74,512 total observations.

Analysis focused on non-Hispanic whites because this is the population in which the wellbeing GWAS was conducted and it is uncertain if a polygenic score derived from this GWAS will show comparable performance in populations with different ancestry (22–24). Because there are many fewer spousal pairs in the genetically-informed sample of non-white HRS respondents, we could not perform parallel analysis of the non-white sample.

## Measures

### Wellbeing Polygenic Score

We calculated HRS respondents' wellbeing polygenic scores based on the Social Science Genetic Association Consortium (SSGAC) GWAS of subjective wellbeing (16). The original GWAS included HRS respondents. To avoid upward bias to polygenic-score-wellbeing associations, SSGAC provided GWAS weights estimated from data excluding the HRS. Therefore, there was no overlap between the samples used to develop the polygenic score and the sample we analyzed. Polygenic scoring was conducted according to the methods described by Dudbridge (25) following the protocol used in previous work (22,26,27). Briefly, we matched SNPs in the HRS genotype database with wellbeing GWAS SNPs. For each SNP, counts of wellbeing-associated alleles were weighted by GWAS-estimated effect-sizes. Weighted counts were summed across SNPs to compute polygenic scores.

To account for any population stratification in our non-Hispanic white sample that might bias polygenic score-wellbeing associations, we computed residualized polygenic scores (26,28). Population stratification is the non-random patterning of allele frequencies across ancestry groups. To quantify any such patterning, we estimated principal components from the genome-wide SNP data for non-Hispanic white HRS respondents according to the method described by Price and colleagues (29) using the PLINK command pca (30). We then residualized the wellbeing polygenic score for the first 10 principal components, i.e. we regressed HRS respondents’ polygenic scores on the 10 principal-component scores and computed residual values from the predictions. Residualized polygenic scores were standardized to M=0, SD=1 for analysis.

### Death of Spouse

Death of one's spouse is a well-documented risk factor for developing depressive symptoms (31). During follow-up, n=8,070 (85%) of the sample were married and n=1,829 (20% of the sample, 24% of those married) experienced the death of their spouse. These survivors (74% female, mean age 73 years) completed follow-up surveys on average 12 months after their spouse's death (IQR=6-18 months, **Supplemental Figure 1**).

### Depressive Symptoms

Depressive symptoms were measured with the 8-item Centre for Epidemiological Studies Depression (CESD) scale at each HRS follow-up. The CESD is a valid psychometric instrument for assessment of depression in older adults (32,33). We analyzed the continuous CESD score. There was substantial within-person variability in depressive symptoms across follow-up (ICC=0.48, M=1.2, SD=1.8).

## Analysis

We conducted descriptive analysis of HRS respondents' depressive symptoms before and after the death of their spouses using non-parametric local regression. Local regression fits a smoothed curve to data that include nonlinear patterns by analyzing “localized” subsets of the data to build up a function iteratively (34). We used local regression to fit symptom data to time-from-death. The resulting model describes symptom levels leading up to and following spousal death. If the death of one's spouse produces a change in depressive symptoms, this change will appear as a discontinuity in the local regression line estimated by the model, e.g. depressive symptom levels may “jump” from their trajectory before the death to a higher level after the death.

We conducted hypothesis tests using parametric nonlinear regression (35). Nonlinear regression estimates a function describing non-linear patterns in a set of data, and then models those nonlinearities as a function of other variables. Analysis modeled depressive symptoms at the first assessment following death of a spouse as a non-linear function of time since the death. We tested the hypothesis that wellbeing polygenic score would buffer against increase in depressive symptoms following death of a spouse using the coefficient for the association between polygenic score and that non-linear function.

Specifically, we estimated the model

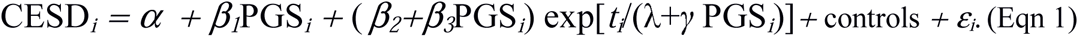

*α* is the model intercept. *ε_i_* is a normally distributed random error term. *β_1_* estimates the main effect of the polygenic score on depressive symptoms. 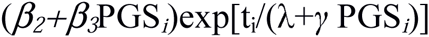 estimates the dynamics of depression in the wake of spousal death. *t*_*i*_ is time since the death, which is >0 for all observations. λ+γ PGS_i_ is the “decay function” for the shock caused by the death. The decay function describes how the increase in depressive symptoms following death attenuates over time. The decay function has two components. λ is the population average decay function. *γ* describes how the decay function varies according to polygenic score. *β_2_* estimates the main effect of spousal death. *β_3_* tests the hypothesis that higher wellbeing polygenic score buffers against increases in depressive symptoms following spousal death. If *β_3_*<0, we reject the null hypothesis that polygenic score is unrelated to depressive symptoms following spousal death. Models included controls for birth year, age, and sex. Graphical representation of model components is shown in **Supplemental Figure 2**.

Statistical analysis was conducted using the R software (36). We tested main-effect associations between HRS respondents’ polygenic scores and their depressive symptoms using random-effects models implemented with the R package lme4 (37). We fit local regression models to repeated measures depressive symptom data using the loess function and the nonlinear regression models using the nls function.

## RESULTS

### Higher wellbeing polygenic score was associated with reduced depressive symptoms

HRS respondents with higher polygenic scores experienced fewer depressive symptoms during follow-up (b=−0.11, CI= −0.14, −0.08, Figure 1). This effect-size is consistent with previous analysis of the wellbeing polygenic score and depression (16). The genetic association with depressive symptoms was similar for men and women (p=0.37 for test of interaction between polygenic score and sex).

**Figure 1.**
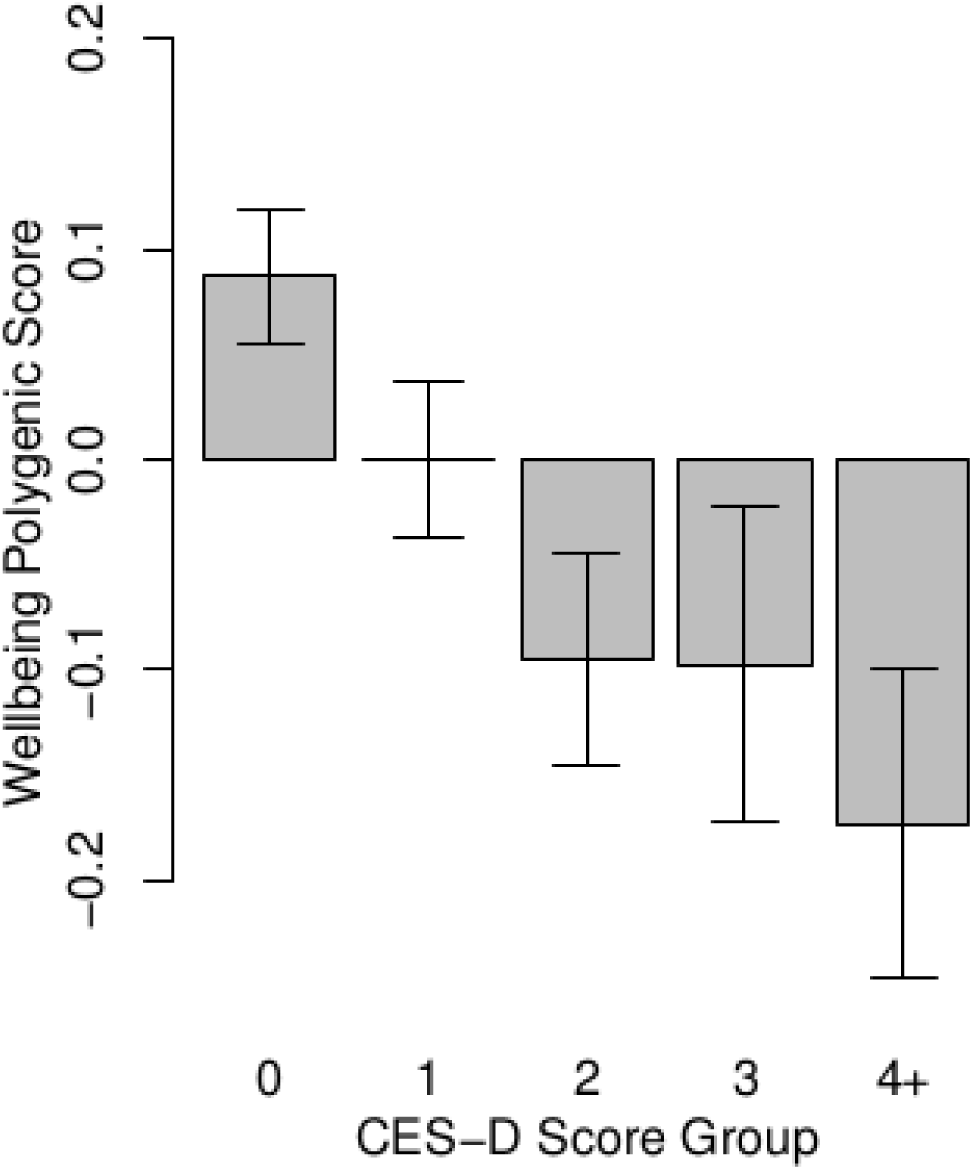
HRS respondents with higher polygenic scores reported fewer depressive symptoms during follow-up. The figure shows average polygenic scores (denominated in standard deviation units) for groups of HRS respondents defined by their histories of depressive symptoms. Depressive symptom history was measured by averaging respondents' CESD scores across repeated assessments and rounded to the nearest integer value. The resulting groups included 3,757 respondents with an average score of 0, 2,819 with a score of 1, 1,470 with a score of 2, 702 with a score of 3, and 705 with a score of 4 or higher.

### Wellbeing polygenic score was not associated with likelihood of experiencing death of a spouse

Genetic liability to depression may influence exposure to stressful life events (38). Therefore, before testing the genetic buffering hypothesis, we evaluated possible gene-environment correlation between subjective wellbeing polygenic score and likelihood of experiencing spousal death. We found no evidence of such gene-environment correlation; HRS respondents' polygenic scores were not related to their likelihood of experiencing the death of their spouse (OR=0.96, CI=0.92−1.01, p=0.145).

### Death of a spouse was associated with an immediate increase in depressive symptoms, followed by a gradual return toward baseline over the following years

HRS respondents who experienced death of their spouse experienced a discontinuity in depressive symptoms around the time of the death. In local regression analysis, respondents' depressive symptoms trended slightly upwards in the months immediately preceding the death, rose sharply at the time of death, and gradually returned toward baseline over the following two years (Figure 2). This pattern of is consistent with previous reports on the course of depressive symptoms around spousal death (39).

**Figure 2.**
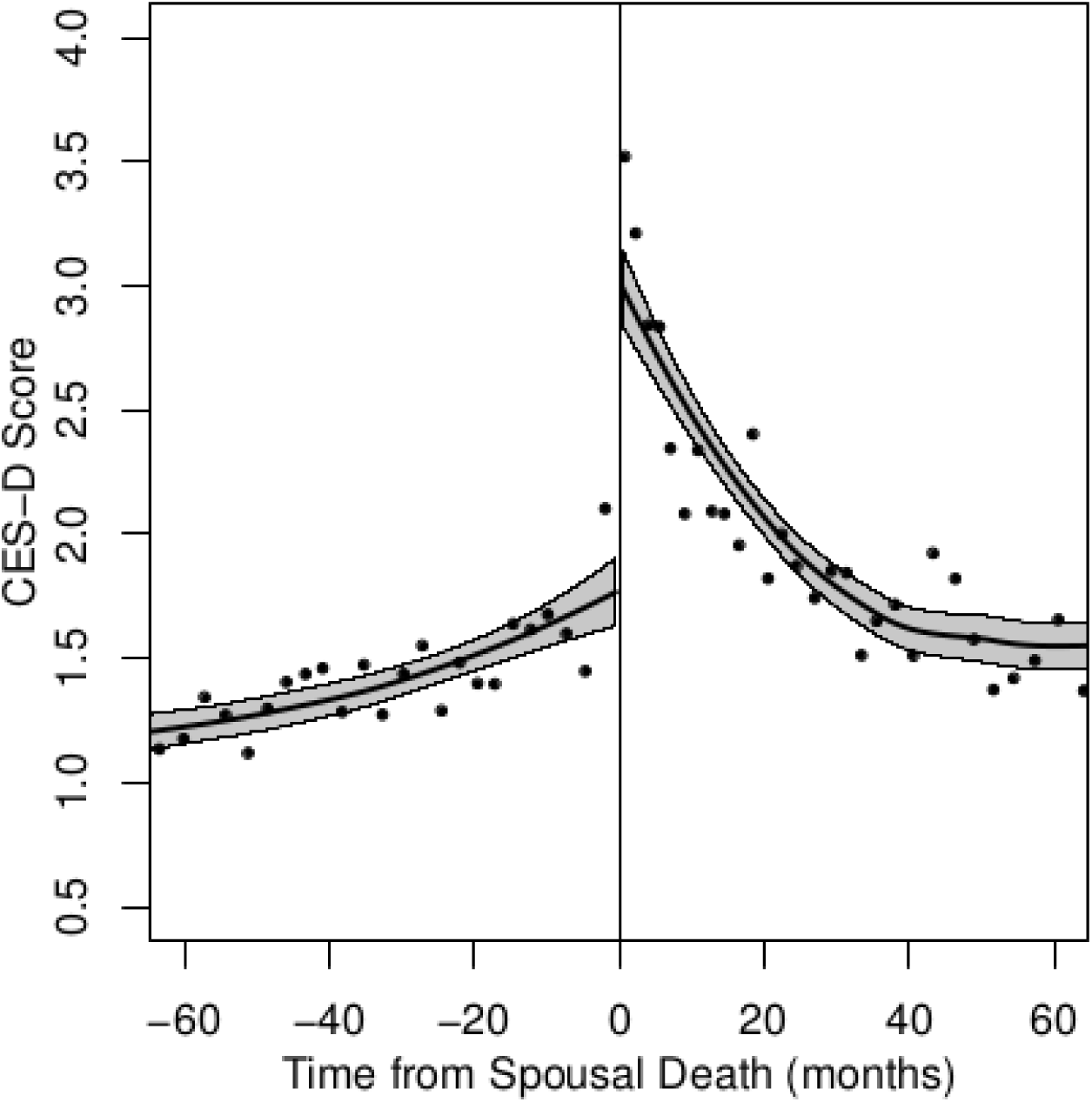
Trajectories of depressive symptoms in the months surrounding the death of a spouse. The figure shows local regression plots of depressive symptoms (CESD scores) by month of measurement relative to the death of a spouse. The plot reflects data from HRS respondents who experience spousal death (N=1,829; 15,853 total observations). Plotted points show mean x- and y- coordinates for bins of about 40 individuals.

Nonlinear regression analysis estimated the increase in CESD associated with spousal death to be 1.9 points (95% CI 1.5-2.3). This increase was smaller for respondents interviewed farther from the time of their spouse's death. Based on trajectories estimated from the model, the increase in depressive symptoms that occurred following the death of a spouse attenuated by 12 months after the death, with modest further attenuation through 24 months (**Supplemental Table 1)**.

### Genetics of subjective wellbeing buffered against increase in depressive symptoms following death of a spouse

HRS respondents with higher polygenic scores experienced flatter trajectories of depressive symptoms around the time of their spouse's death as compared to peers who also lost their spouse and had lower polygenic scores. In local regression analysis, HRS respondents with higher wellbeing polygenic scores experienced fewer depressive symptoms in the months leading up to the death and a smaller increase in depressive symptoms following the death as compared to other survivors who had lower polygenic scores (Figure 3).

**Figure 3.**
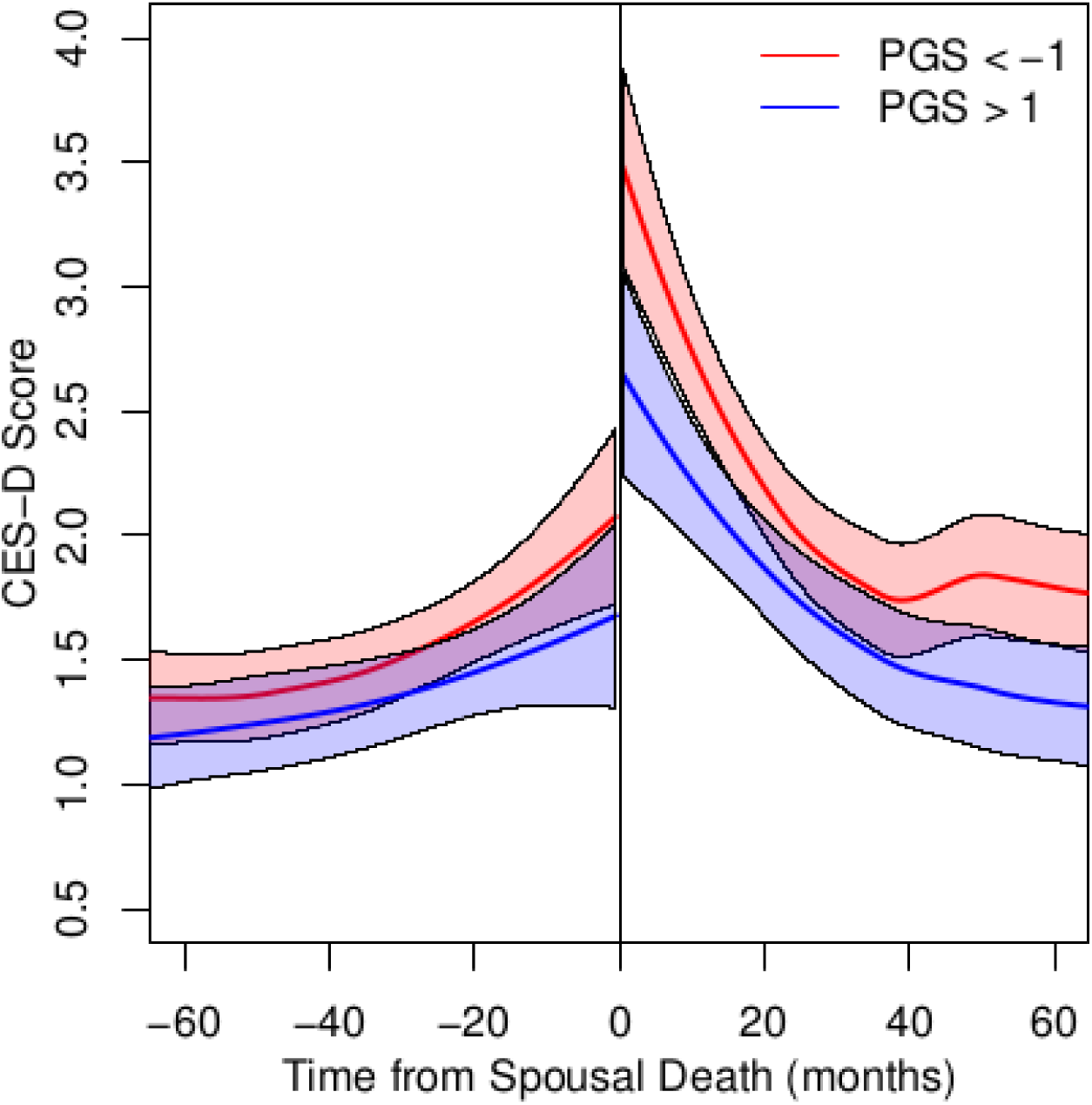
Trajectories of depressive symptoms for HRS respondents with low polygenic scores (red line) and high polygenic scores (blue line). The figure shows local regression plots of depressive symptoms (CESD Scores) by month of measurement relative to the death of a spouse. Trajectories for respondents with low polygenic scores (1 or more SDs below the mean, n=2,493) are graphed in red. Trajectories for HRS respondents with high polygenic scores (1 or more SDs above the mean, n=2,181) are graphed in blue.

In nonlinear regression analysis, each standard-deviation increase in polygenic score buffered the increase in depressive symptoms following spousal death by 0.46 CESD points (95% CI 0.06, 0.86, p=0.02). In context, this effect estimate suggests a person with a polygenic score 1SD above the population mean should experience about half as much increase in depressive symptoms following death of their spouse as compared to a peer whose polygenic score was 1 SD below the population mean. Respondents' polygenic scores were not related to the rate of attenuation in depressive symptoms following the death (γ=0.41, p=0.72, **Supplemental Figure 3, Supplemental Table 1**).

### Sensitivity Analyses

We conducted sensitivity analyses to evaluate consistency of findings. First, we repeated analysis in a subset of HRS respondents identified as being of European descent based on analysis of their genetic data (N=1,647). (Our original analysis included all respondents who self-identified as non-Hispanic white.) Findings were unchanged in the genetically-defined European-descent sample (**Supplemental Table 2**).

Second, we repeated analysis using polygenic scores derived from two published GWAS of depression, the Psychiatric Genomics Consortium's GWAS of major depressive disorder (40) and Social Science Genetic Association Consortium GWAS of depressive symptoms (16). The subjective wellbeing score was negatively correlated with depression polygenic scores (r=−0.16 with the major depression score and r=−0.32 with the depressive symptoms score; the major depression score and depressive symptom scores were correlated with each other, r=0.28). Results from analysis of depression polygenic scores paralleled results from analysis of the subjective wellbeing polygenic score, but were in the opposite direction, reflecting the negative correlation between depression polygenic scores and subjective wellbeing polygenic score. HRS respondents with higher depression polygenic scores had elevated baseline levels of depressive symptoms and experienced a larger increase in depressive symptoms following the death of a spouse as compared to respondents with lower depression polygenic scores (**Supplementary Table 3**).

## DISCUSSION

We tested if genetics discovered in GWAS of subjective wellbeing buffered against development of depressive symptoms following death of a spouse. We analyzed data on depressive symptoms in 9,453 individuals followed longitudinally as part of the US Health and Retirement Study (HRS). During follow-up, 1,829 of these individuals experienced the death of their spouse. HRS respondents with higher polygenic scores experienced fewer depressive symptoms during follow-up. For those whose spouse died during the follow-up period, having higher polygenic score buffered against increased depressive symptoms following the death. Having a low polygenic score for depression had similar buffering effects on depression risk to having a high polygenic score for subjective wellbeing. Magnitudes of genetic effects were small.

This evidence of genetic buffering against depression following a stressful life event provides proof of concept for further investigation of the genetics of subjective wellbeing as a modifier of the life-stress to depression link. Studies of gene-environment interplay are controversial (41). Statistical and theoretical models are contested (42–45). The precise nature of any “interaction” between the stress of losing a spouse and wellbeing-related genetic background remains uncertain. What we find is that individuals who carry more wellbeing-associated alleles tend to develop fewer additional depressive symptoms following the death of their spouse. This “buffering” is independent of the overall lower level of depressive symptoms associated with wellbeing polygenic score in the absence of a spousal death. Buffering is also not affected by any “gene-environment correlation” between genetics of subjective wellbeing and exposure to death of a spouse (46); HRS respondents’ subjective wellbeing polygenic scores were unrelated to their risk of losing a spouse during follow-up. Finally, we detected consistent evidence of buffering using two different statistical specifications, local regression and nonlinear regression, that take into account time elapsed between the stressful event and the measurement of depressive symptoms.

Findings have implications for relevance of wellbeing genetics in clinical approaches to bereavement and for research into gene-environment interplay. Clinically, genetic testing based on current knowledge is unlikely to be useful in planning for or managing bereavement. Effect sizes in our study were too small to be useful for predicting outcomes of individual patients. It is possible that larger-scale GWAS will furnish more predictive polygenic scores with greater clinical relevance. However, even with very-large GWAS, predictive power of polygenic scores for common, complex traits will be limited (47). Instead, our results may inform clinical research into candidate intervention targets to promote resilience in bereavement and in the context of other stressful experiences. Specifically, our findings raise questions about what behaviors or psychological processes mediate buffering effects. Do individuals with higher wellbeing polygenic scores process or cope with stress differently (48)? Do they cultivate different social support networks or interact with them in different ways? Studies to identify such mediators of genetic associations can inform interventions designed to promote resilience (49).

Regarding research on gene environment interplay, our results highlight two issues related to study design. First, our study suggests a model for integrating polygenic scores into studies of gene environment interplay: consider polygenic scores for characteristics theorized to modify effects of environmental exposures. Polygenic scores accumulate information from across the genome and take on normal distributions—properties that match theoretical models of human individual differences (50). But they are derived from GWAS that are naïve to environmental variation. Consequently, polygenic scores may quantify precisely those genetic influences that do not vary depending on environment as a byproduct of standard GWAS study design, although alternative designs are possible (51,52). If correct, this explanation could account for null findings in previous studies of gene-environment interplay in depression that used polygenic scores derived from depression GWAS (9,10). In contrast, we studied a polygenic score for a characteristic theorized to buffer environmental risks for depression. This “triangulation” approach may prove useful to future investigations. That said, our analysis of polygenic scores for depression yielded substantively identical results to our analysis of the polygenic score for subjective wellbeing. Thus, the observed gene-environment interplay is also consistent with a diathesis-stress model. Further investigation of overlap between genetic underpinnings of subjective wellbeing and depression in the context of stressful life events is warranted.

A second design issue raised by our study is the temporal resolution of environmental exposure measures. Depression is an episodic condition. Even persistent cases experience temporary remissions. Increases in depressive symptoms following stressful life events may attenuate over time, as is characteristic in bereavement (53). Our study could account for this attenuation because precise dates were available for the timing of spousal death. We used statistical models designed for this type of data. Future studies may also wish to focus on stressful life events that can be located in specific temporal relation to the measurement of depressive symptoms (54).

This study has limitations. We studied depression in married older adults. Etiology of depression in older adults may differ in some ways from etiology in younger adults (55), although the relationship of stressful life events to depression does not seem to vary with age (56). Etiological features of depression, including subjective wellbeing may influence probability of marriage and of remaining married into later life (57). Replication of results in younger samples and with unmarried adults is needed. We studied depression following death of a spouse. Depression following death of a spouse may be different from depression related to other stressful life events. However, the distinction is unclear, as is reflected in the most recent revision of the American Psychiatric Association's Diagnostic and Statistical Manual of Mental Disorders, 5th Edition (58). There is evidence that so-called bereavement-related depression shares etiology with depression related to other stressful life events (59). Tests of wellbeing polygenic score as a buffer against depressive symptoms following other types of stressful life events are needed. Our analysis was restricted to European-descent HRS participants. We applied this restriction because of concerns about generalizability of GWAS results across racial/ethnic populations (24) and because the sample of spousal pairs of non-white HRS participants was too small for an independent analysis. Although relationships among stress, coping, and depression may be similar across ethnic groups (60), studies of the genetics of wellbeing and their potential role in buffering against depression following stressful life events in non-European populations are needed.

Research into interplay between genes and environments has a contentious history in psychiatry. With the advent of large-scale GWAS, new, more reliable genetic measures of liability to psychiatric disorders and other mental health characteristics are emerging. At the same time, genetic data are becoming available for large, population-based social surveys that have recorded environmental exposure information for participants. Together, these resources should enable a new generation of molecular genetic research into gene-environment interplay research in the etiology of psychiatric disorders.

## Acknowledgements

This research uses data from the HRS, which is sponsored by the National Institute on Aging (Grants NIA U01AG009740, RC2AG036495, and RC4AG039029) and conducted by the University of Michigan. Further support was provided by the NIH/NICHD-funded University of Colorado Population Center (R24HD066613). DWB is supported by an Early Career Research Fellowship from the Jacobs Foundation and NIA grants P30AG034424 and P30AG028716. AO is supported by ERC Consolidator Grant (647648 EdGe). This research was facilitated by the Social Science Genetic Association Consortium (SSGAC).

